# How much data do we need? Reliability and data deficiency in global vertebrate biodiversity trends

**DOI:** 10.1101/2023.03.18.532273

**Authors:** Shawn Dove, Monika Böhm, Robin Freeman, Louise McRae, David J. Murrell

**Affiliations:** Centre for Biodiversity and Environment Research, University College London, Gower Street, London WC1E 6BT, UK; Institute of Zoology, Zoological Society of London, Outer Cir, London NW1 4RY, UK; Global Center for Species Survival, Indianapolis Zoo, 1200 West Washington Street, Indianapolis, IN 46222, USA

**Keywords:** Living Planet Index, LPI, biodiversity indicators, biodiversity trends, biodiversity data, data deficiency, global biodiversity monitoring, indicator testing, indicator accuracy, indicator reliability

## Abstract

Global biodiversity is facing a crisis, which must be solved through effective policies and on-the-ground conservation. But governments, NGOs, and scientists need reliable indicators to guide research, conservation actions, and policy decisions. Developing reliable indicators is challenging because the data underlying those tools is incomplete and biased. For example, the Living Planet Index tracks the changing status of global vertebrate biodiversity, but taxonomic, geographic and temporal gaps and biases are present in the aggregated data used to calculate trends. But without a basis for real-world comparison, there is no way to directly assess an indicator’s accuracy or reliability. Instead, a modelling approach can be used.

We developed a model of trend reliability, using simulated datasets as stand-ins for the "real world", degraded samples as stand-ins for indicator datasets (e.g. the Living Planet Database), and a distance measure to quantify reliability by comparing sampled to unsampled trends. The model revealed that the proportion of species represented in the database is not always indicative of trend reliability. Important factors are the number and length of time series, as well as their mean growth rates and variance in their growth rates, both within and between time series. We found that many trends in the Living Planet Index need more data to be considered reliable, particularly trends across the global south. In general, bird trends are the most reliable, while reptile and amphibian trends are most in need of additional data. We simulated three different solutions for reducing data deficiency, and found that collating existing data (where available) is the most efficient way to improve trend reliability, and that revisiting previously-studied populations is a quick and efficient way to improve trend reliability until new long-term studies can be completed and made available.

## 1. Introduction

An urgent data crisis complicates the global biodiversity crisis (Turak et al., 2017). Attempts to assess global biodiversity (e.g. the Intergovernmental Science-Policy Platform on Biodiversity and Ecosystem Services, IPBES) and to set global policies and goals that will halt or reverse its loss (e.g. the Convention on Biological Diversity, CBD, and Sustainable Development Goals, SDGs) need reliable and up-to-date scientific information (Jetz et al., 2019). Yet most studies and tracking programs are either species- or region-focused, temporally limited and inherently biased, all of which results in large geographic and taxonomic knowledge gaps (Hortal et al., 2015; Jetz et al., 2019; Meyer et al., 2015; Proença et al., 2017; Turak et al., 2017). Advances in technologies such as camera tracking, satellite sensors, digital image recognition, network speed and capacity, data access, and mobile devices are improving our ability to track and count populations of birds and mammals (Lausch et al., 2016; Nichols et al., 2011; Rose et al., 2015), but our datasets are far from complete. The situation is worse for amphibians, reptiles, insects, and other groups, for which many species have yet to even be described (Mora et al., 2011).

We need tools to improve our understanding of global biodiversity within the limitations imposed by biased and incomplete datasets. Mace & Baillie (2007) suggested a solution: develop indicators based on existing data, understand data biases, and develop methods to reduce the bias. Biodiversity indicators summarize complex scientific information in a simple way, often serving as a bridge between science and policy (Secretariat of the Convention on Biological Diversity, 2006). But what can we expect from indicators that summarize only a fraction of the biodiversity they represent? To what extent can we rely on them to present a true picture of the state of global biodiversity?

Two of the best-known biodiversity indicators are the Living Planet Index (LPI), which tracks vertebrate population trends (McRae et al., 2017), and the Red List Index (RLI), which tracks extinction risk trends (Butchart et al., 2005). The RLI is based on extinction risk classifications at the species-level, created by expert assessment using an objective set of criteria (IUCN, 2012). By contrast, the LPI uses continuous population data collected by scientific surveys. However, as intensive global long-term studies do not exist for most species, the LPI calculates trends from data compiled from a variety of sources, including grey literature (McRae et al., 2017). This means a lack of standardization in study design (individual population time series are standardized, but there is no standardization between populations), monitoring strategy, frequency of assessment, monitoring intensity and effort, even data type (densities, counts of individuals or breeding pairs or even nests, and population size estimates are mixed together). The LPI has taxonomic and geographical imbalances (Collen et al., 2009; McRae et al., 2017), a problem found also in other global biodiversity datasets (Boakes et al., 2010; Collen et al., 2008; Yesson et al., 2007). Further, many included time series are short (McRae et al., 2016; Proença et al., 2017; Saha et al., 2018), and shorter trends tend to be less accurate than longer ones (Arkilanian et al., 2020; Wauchope et al., 2019). Recognizing these limitations, the LPI employs statistical techniques to increase the accuracy and precision of trends. Generalized Additive Modelling or log-linear interpolation are used (depending on the length of a given time series) to fill in missing values in time series, bootstrapping is used to generate confidence intervals (Collen et al., 2009), and a hierarchical weighting system is applied to account for geographical and taxonomic bias (Collen et al., 2009; McRae et al., 2017).

Without a basis for real-world comparison, there is no way to directly assess an indicator’s accuracy or reliability. However, there are ways to address this question indirectly. One solution was employed by the sampled approach to the Red List Index (sRLI), which uses the minimum representative sample size needed to achieve less than a 5% probability of falsely detecting a positive slope when the Red List Index trend is negative (Baillie et al., 2008; Henriques et al., 2020). Minimum representative sample size was determined through sub-sampling of comprehensively assessed species groups on the IUCN Red List (e.g. mammals, birds etc.; Baillie et al., 2008; Henriques et al., 2020).

Two challenges presented by the LPI require a different approach than that taken for the sRLI. First, LPI trends are based on population time series that are often short and/or infrequently measured, and there are no regional or taxonomic groups within the LPI where the data is comprehensive enough to be certain of the real-world trend. Therefore, comparing sampled trends to LPI trends would tell us little about how the sampled trends might compare to reality. Second, the LPI uses non-linear trends that change slope and direction over time, so trends should be compared in a way that reflects this. Here, we use a modelling approach to overcome these challenges, based on thousands of datasets of synthetic population time series with variations in the underlying properties of the data to represent regional taxonomic groups in the real world and sampling from those datasets. We degraded the samples by randomly removing observations and adding observation error to resemble regional taxonomic groups in the Living Planet Database (LPD, the database underlying the LPI). We then compared the trends calculated from the samples with those from the complete datasets using a distance metric (Dove et al., 2022), and constructed a multiple regression model to understand how the distance values are influenced by variations in properties of the data. Here, distance metrics can be thought of as a measure of trend accuracy. By selecting a threshold value for accuracy and applying the model to the LPI, we were able to quantify the reliability of disaggregated LPI trends and determine the number of additional time series needed to meet the threshold. Finally, we modelled and compared three different solutions for reducing data deficiency: a) tracking unstudied populations for a decade to generate new time series for the LPD, b) resampling previously-studied populations to update old time series in the LPD, and c) gathering more time series from existing studies to add to the LPD. The results from this study can be used to focus data-gathering and data-collation efforts on the regions, taxa, and populations that would be of greatest benefit to improving our understanding of the state of global vertebrate biodiversity.

## 2. Material and Methods

Fig. 1 shows an overview of our methods, with each numbered step corresponding to a numbered subheading in the text.

**Fig. 1.**
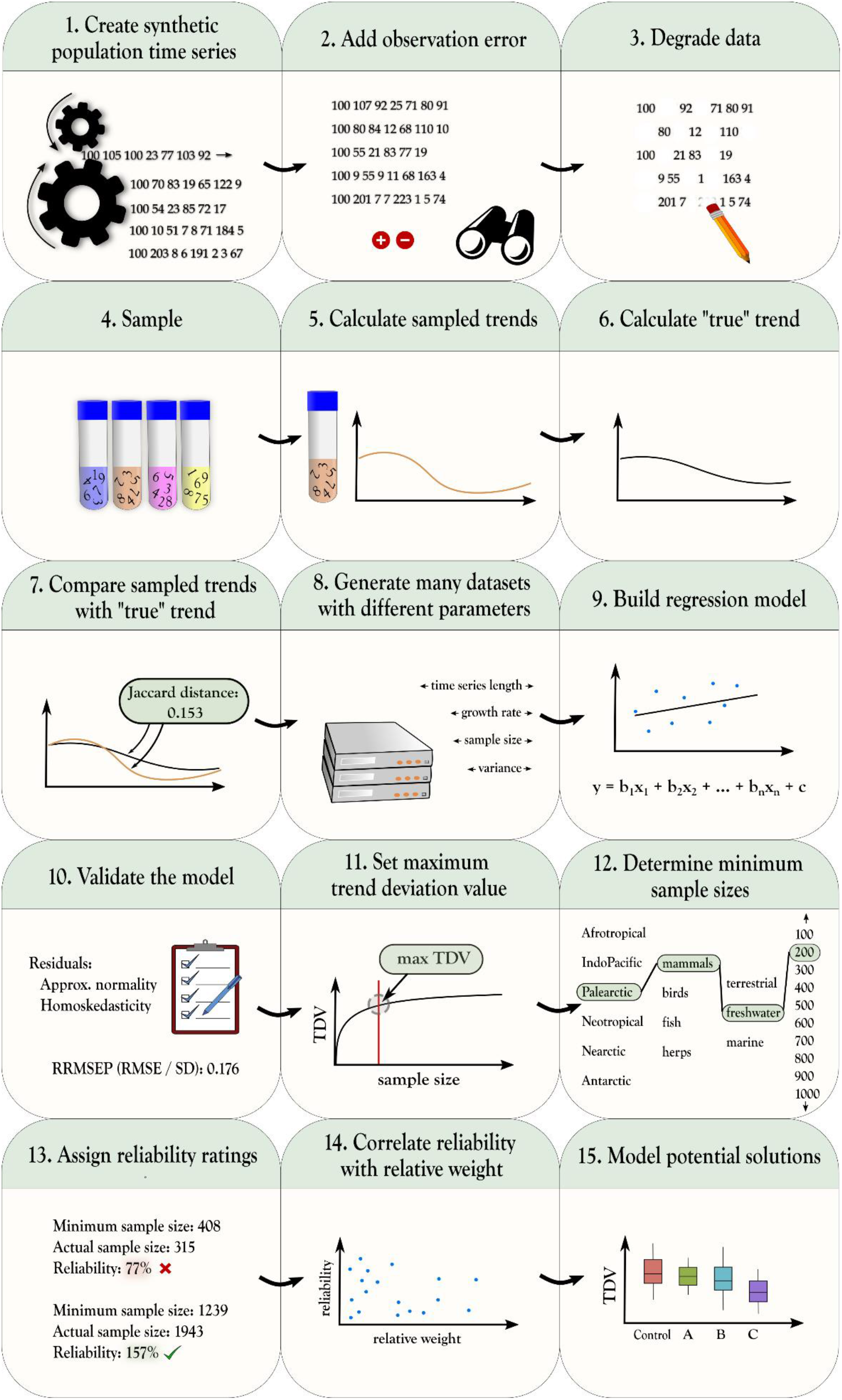
Modelling trend accuracy in the LPI: an overview.

### 2.1. Synthetic data generation

We first created simulated datasets to represent “real-world” regional vertebrate groups for which the LPI calculates biodiversity trends. The LPI is often represented as a single global index trend, but can also be disaggregated into hierarchical groups: first into systems (terrestrial, marine, freshwater), then geographical realms within each system, and finally taxonomic groups within each realm. It is this lowest level of the hierarchy, the regional taxonomic groups, which we simulated. From here on simulated regional taxonomic groups will be referred to as datasets. The base units of the LPI, and of our synthetic datasets, are population time series, which we will refer to simply as populations. These populations are grouped into species, and species are grouped into datasets.

Our procedure to simulate a dataset requires six parameters: 1) the total number of populations to simulate (set to 10,000), 2) the mean number of populations assigned to each species (set to 10), 3) the number of years (length of trend) to simulate (set to 50), 4) the mean of the population mean growth rates (*μ_ds_*), 5) the standard deviation of the population mean growth rates (variation among populations, *σds*), and 6) the mean of the population standard deviations of the growth rate (process error, *μ_ɳ_*). The first three parameters were fixed. The first, total populations, affects trend accuracy only when greater than half of all populations in a dataset are sampled (see Fig. S1), a situation that is unlikely for regional taxonomic groups in the LPD, as it is rare even at the species level (see taxonomic representativeness in McRae et al., 2017). The second parameter, the mean number of populations per species, has no effect on trend accuracy within the wide range of values we tested (see Fig. S2). The third, trend length, is constant across regional taxonomic groups in the LPD. However, it does affect trend accuracy (see Fig. S3), and would therefore need to be set appropriately if adapting the model for a different indicator. Parameters four through six are variable in the LPD and affect trend accuracy, and were therefore set to vary in the simulations.

We model population time series using the stochastic exponential model with process error:

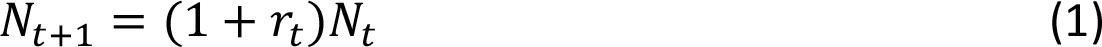

where *N_t_* is population size at year *t*, 1 + *r_t_* is annual growth rate at year *t*, and 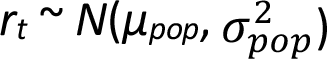 models uncorrelated process error (i.e. temporal variation in the growth rate that could be caused by, for example, uncorrelated environmental variation) by sampling each annual growth rate from a normal distribution. Process error, *ɳ*, is represented by the mean of the population standard deviations of *r*, with 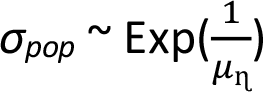.

The mean of the normal distribution of population growth rates is itself drawn from a normal distribution, 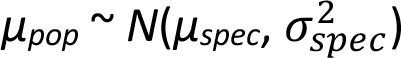. Thus, populations from a species will have similar but not identical underlying mean population growth rates representing perhaps differences in environmental conditions between geographically isolated populations of a given species. In turn, similar species are grouped together into datasets, and we assume that species within taxonomic groups have underlying population growth rates that are drawn from an identical distribution, 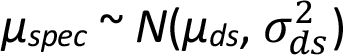.

Here, larger values for *σ*_*ds*_lead to a broader range of underlying species growth rates, perhaps signifying broader species-specific variation in responses to drivers such as habitat change within a taxonomic group. Using this hierarchical approach therefore captures the similarity of time series within a species, and the similarity of time series between species within a taxonomic group.

Growth for each population was modelled for 50 years, starting at a population size of 100. Populations were assigned to species by randomly sampling from a pool of 500 species IDs, with replacement, resulting in a normal distribution of populations per species, 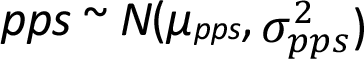, with *μ_pps_* = 20 and *σ_pps_* = 4.5. While populations are unlikely to be normally distributed across species in the real world (one would expect more rare species than common species), simulations confirmed that our modelling approach is robust against distributional assumptions for this parameter (see Fig. S2).

### 2.2. Observation error

The variation in lambdas modelled above assumes all variation is due to process error. However, time series in the LPD are based on population estimates, which can be assumed to include some level of observation error due to e.g. species misidentification, non-detection, and counting errors. This observation error is not accounted for in the LPI, but may affect trend reliability. Observation error, ɛ, can be calculated using the coefficient of variation (cv), defined as

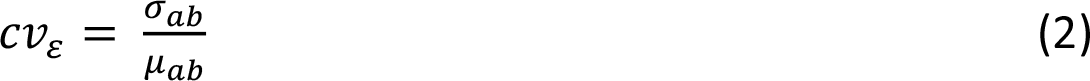

where *μ_ab_* and *σ_ab_* are the mean and standard deviation (respectively) of the abundance values. Since data in the LPD was collected using a variety of methods, and ɛ is not recorded in the database, we chose a range of *ɛ* consistent with values reported for other vertebrate surveys (Fryxell et al., 2014; Westcott et al., 2012; Zylstra et al., 2010). We determined through simulations that there is no effect of increasing observation error on trend accuracy (Fig. S4), therefore an approximate range of *ɛ* should suffice. For each simulated population time series, *ɛ* was randomly selected from a normal distribution with *μ_ɛ_* = 0.15 and *σ_ɛ_ = 0.1*. We modelled observed abundances of a population at a time point, *Z*_*t*_, as

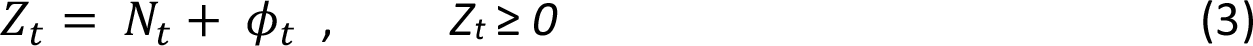

where *N_t_* is the population size at time *t* taken from equation (1) and *ϕ_t_* is a normally distributed variable, 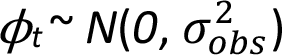, with

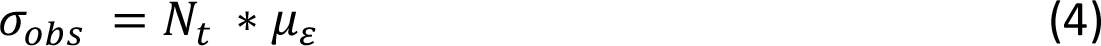

where *σ_obs_* is the standard deviation of *ϕ_t_*. A value for *μ_ɛ_* of 0.1 (10%) would result in approximately 68.2% of observations falling within 10% of their corresponding simulated ‘true’ values and 99.7% of simulated observations falling within 30%.

### 2.3. Data degradation

Observed versions of the datasets were then randomly degraded to resemble the varied quality of sampled real-world data present within the LPD. The length (number of years from first to final observation) for each degraded time series within a dataset was randomly chosen by sampling from a Poisson distribution. We determined through simulations that varying the number of observations does not affect trend accuracy at a given time series length, so we fixed the mean number of observations at half of the mean time series length (rounded up). The starting years for each time series were assigned randomly. Time series were then cut to their assigned length, and half of the remaining observations were randomly removed.

### 2.4. Sampling

Populations were randomly sampled from each dataset, without replacement. This was repeated to obtain 20 random samples of the same size for each dataset. Values for four of the six dataset parameters described in Section 2.1 may be different for samples than for the dataset they are selected from, and may also vary between samples: the mean number of populations per species, the mean and standard deviation of population mean growth rates, *μ_ds_* and *σ_ds_*, and the mean of population standard deviations of the growth rate, *μ_ɳ_*.

### 2.5. Calculation of sampled trends

Non-linear index trends were calculated from each sample, following the LPI method described in McRae et al. (2017). First, time series with six or more data points were modelled using a Generalized Additive Model (GAM), as described in Collen et al. (2009), with a Gaussian (normal) distribution, smoothed by a thin plate regression spline, with the number of knots set to half the number of observations (rounded down). The model fit was checked by applying a GAM to the residuals, this time smoothed by a shrinkage version of a cubic regression spline, with the number of knots set to the full number of observations of the time series (before the GAM model was applied) and gamma set to 1.4. If the sum of the estimated degrees of freedom from the modelled residuals was close to one (greater than 0.99 and less than 1.01), the population GAM was considered a good fit. Time series that did not pass the model fit test, or that had fewer than six data points, were interpolated using the chain method (Loh et al., 2005), as described in Collen et al. (2009). The chain method imputes missing values using log-linear interpolation by

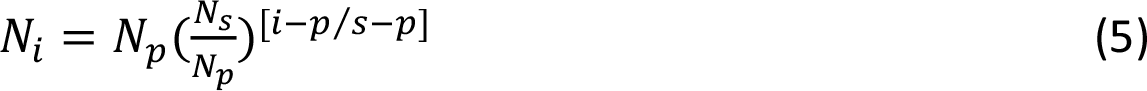

where *N* is the population estimate, *i* is the year for which the value is to be interpolated, *p* is the preceding year with an observed value, and *s* is the subsequent year with an observed value. For all populations, whether interpolated or modelled by a GAM, species indices were formed by a three-step process. First, population sizes were converted to annual rates of change by

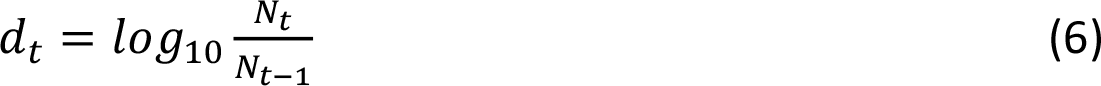

where *N* is the population estimate and *t* is the year. Second, average growth rates were calculated for each species by

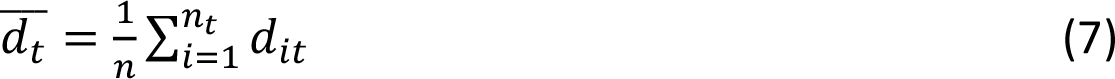

where *n*_*t*_ is the number of populations in a given species, *d*_*it*_ is the growth rate for population *i* at year *t*, and 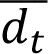 is the average growth rate at year *t*. Growth rates were capped at [-1:1]. Finally, index values were calculated by

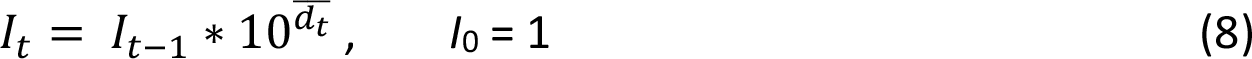

where *I* is the index value and *t* is the year. Equations 5-8 are from Collen et al. (2009).

### 2.6. Calculation of the *‘*true*’* trend

A non-linear index trend was calculated for each complete, undegraded dataset (without observation error), following McRae et al. (2017), as for the sampled trends. However, the undegraded datasets have no missing values, therefore modelling each time series using the chain method or a GAM was unnecessary and that step was skipped.

### 2.7. Comparison of trends

We selected an appropriate distance measure to compare sampled trends with ‘true trends’ using the process described in Dove et al. (2022). Of the distance measures deemed appropriate, we chose the Jaccard distance because it uses a 0-1 scale, making it easier to interpret. The Jaccard distance is calculated as

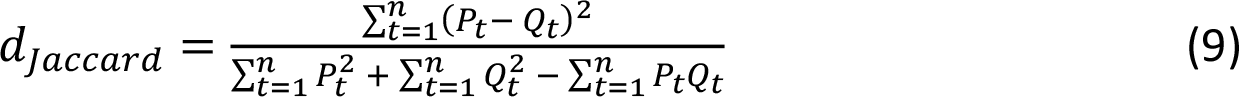

(from Cha, 2007), where P*_t_* and Q*_t_* are index values from two trends P and Q at time point *t*, and *n* is the number of time points. From here on, any value calculated by applying the Jaccard distance to compare sampled versus ‘true’ trends will be referred to as a trend deviation value, or TDV.

We use TDV here as a measure of trend accuracy, but it is in fact the complement of accuracy (a perfectly accurate trend would yield a TDV of zero); lower TDV means higher accuracy. Furthermore, when referring to TDVs of simulated trends, we use the term ‘trend accuracy,’ but when referring to TDVs of LPI trends, we use the term ‘trend reliability.’ This is because TDVs for simulated trends are measured, while TDVs for LPI trends are estimated based on a model. Trend reliability is thus a measure of *expected* accuracy based on underlying data sufficiency or deficiency, but should not be considered a proxy for accuracy. In other words, a data deficient trend may be accurate but we cannot rely on it to be so.

### 2.8. Generation of datasets

We generated 3,000 datasets (each consisting of 1,000 species and 10,000 populations), with each dataset sampled 20 times, resulting in 60,000 samples. Values for mean time series length, *μ_ds_*, *σ_ds_*, and *μ_ɳ_* were randomly selected from uniform distributions, while sample size was randomly selected from a log-uniform distribution, ln(SS) ∼ *U*(ln(a), ln(b)), where SS is sample size and a and b are the minimum and maximum values, respectively (log-uniform was chosen to ensure the model would be robust at small sample sizes, as most datasets in the LPD are small). Ranges for the distributions were chosen to ensure that parameter ranges in the samples would be broader than the ranges present in the LPD (Table 1). Regional taxonomic groups from the LPD with fewer than 20 populations were excluded from parameter range calculations to avoid extreme outliers. We set the minimum sample size to 50 because smaller samples rarely generate a complete trend, and the maximum to 10,000 to improve predictions of the effects of sample size increases.

**Table 1.** Parameters with value ranges for simulated datasets, degraded samples, and the LPD

| Independent Variable | Range in Datasets | Range in Samples | Range in LPD |
| --- | --- | --- | --- |
| Sample Size | – | 50 – 9975 | 2 – 3000 |
| Mean Length of Time Series | 6.0 - 38 | 5.5 – 39 | 6.0 – 39 |
| Mean of Pop. Mean Growth Rates, $\mu_{ds}$ | -0.13 – 0.12 | -0.25 – 0.31 | -0.19 – 0.16 |
| St. Dev. of Pop. Mean Growth Rates, $\sigma_{ds}$ | 0.074 – 0.59 | 0.097 – 0.83 | 0.12 – 0.63 |
| *Mean of Pop. Growth Rate St. Dev., $\mu_{\eta}$ | 0.049 – 1.17 | 0.13 – 1.06 | 0.16 – 0.89 |

\*This parameter is modelled as process error in the simulated datasets, but in the degraded samples it represents process error and observation error combined.

### 2.9. Multiple regression model

We built a multiple linear regression model to understand how variables in the simulated data determine trend accuracy (TDV). First, we removed all simulated datasets in which the mean of the sample parameter values fell outside of LPD parameter ranges (individual replicates were allowed to fall outside of LPD ranges), leaving 2,361 datasets, or 47,220 samples. We then randomly selected 67% of the remaining datasets (1,581 datasets) to train the model. The other 33% (780 datasets) we set aside for testing the model.

### 2.10. Model validation

The residuals of the combined data used to train the model were approximately normally distributed. Likewise, the residuals appeared homoskedastic when plotted against fitted values. We compared the actual TDV of each sample in the testing datasets to the predicted TDV for that sample calculated by the model, then calculated the RRMSEP (relative root mean squared error of prediction), defined as

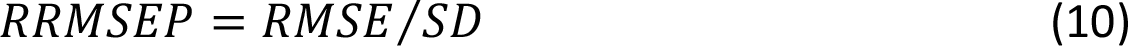

where RMSE is the root mean squared error and SD is the standard deviation of the actual TDVs, and

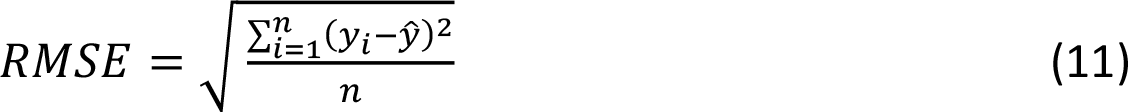

where *y_i_* is the *i*th actual TDV, *ŷ* is the predicted TDV, and *n* is the number of samples.

### 2.11. Maximum trend deviation value

We set a maximum predicted TDV as a threshold that regional taxonomic group trends within the LPI should not exceed to be considered reliable. First, we built a linear regression model of the square root of TDV from our training datasets, with the natural log of sample size as the predictor variable, since sample size is the only user-controlled variable within the LPD. Every regional taxonomic group within the LPD represents a single sample from the real world; therefore, we were not interested in the mean TDV achieved by each dataset, but in the range of possible TDV values, especially the upper part of the range (the least accurate sample trends from each dataset).

We used 10,000 bootstrap estimations of the mean of the TDV from each dataset to calculate the 90% confidence intervals using the bias corrected and accelerated bootstrap interval (BCa) method, also known as the adjusted bootstrap percentile method. The BCa method is a non-parametric method that does not assume the data is normally distributed (the TDV values have a beta distribution) and corrects for bias and skewness in the distribution of the mean estimates. We plotted the curve of the sqrt-log model of the upper 90% confidence interval of TDV in relation to sample size on a (non-log) graph of TDV versus sample size (Fig. 2).

**Fig. 2.**
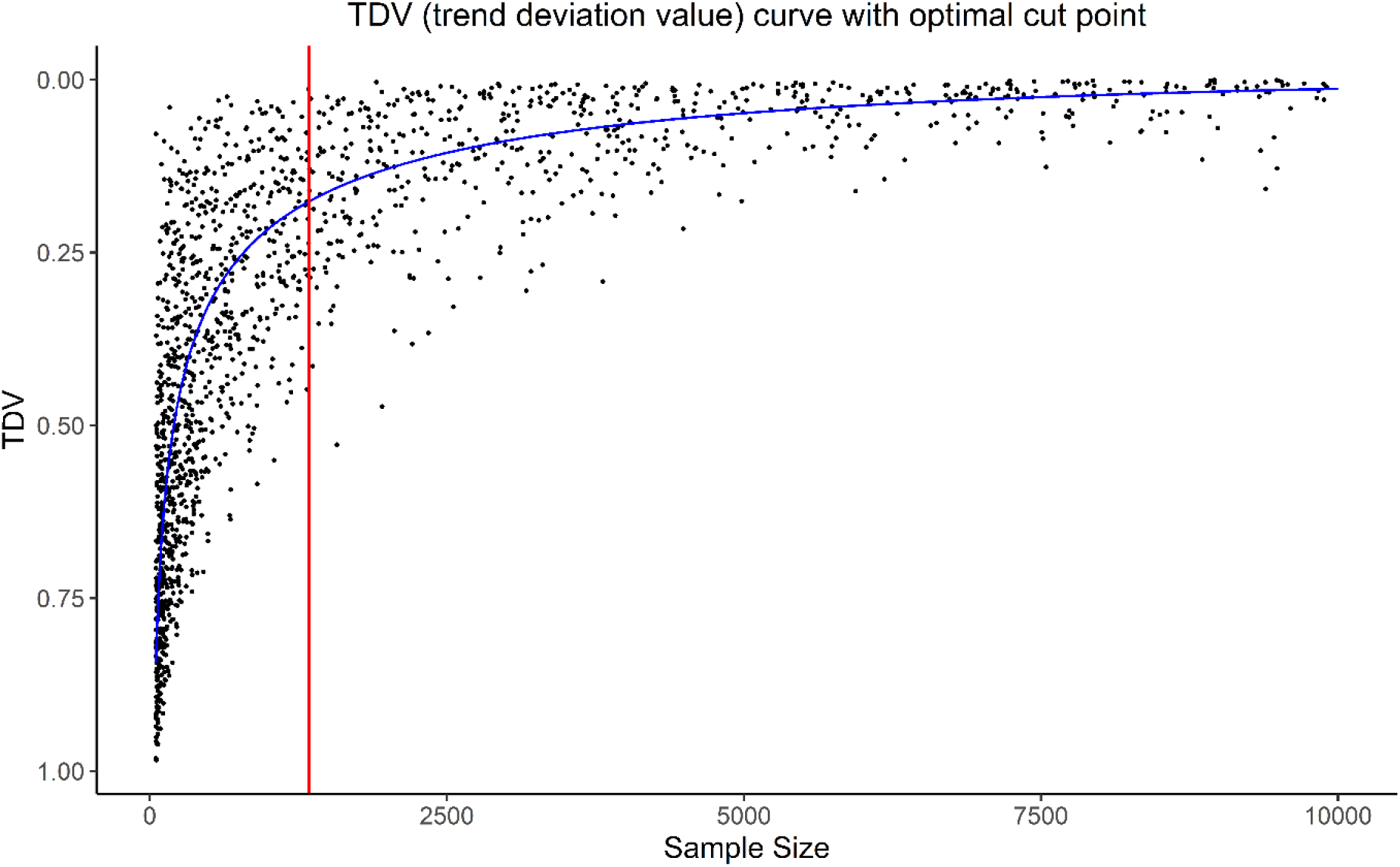
Trend deviation value (TDV) versus sample size. This plot includes only the upper 90% confidence interval of TDV from each simulated dataset. The curved blue line is the sqrt-log model of the plotted values. The vertical red line intersects the sqrt-log curve at the optimal cut-point.

Increasing sample size should naturally lead to more desirable TDV, but it is costly in terms of time and money to increase sample size, and it may also be prudent to put the resources elsewhere. It is therefore important to choose a maximum TDV that reflect these trade-offs. To choose a maximum TDV, we used a method called the concordance probability method (CZ) (Liu, 2012). We adapted CZ from the field of biomedical research, where it is often necessary to specify a cut-off value to discriminate between positive and negative results from screening or diagnostic tests (Liu, 2012). First, a receiver operating characteristic (ROC) curve is built, plotting the rate of true positives (sensitivity) against the rate of false positives (1 – specificity). The idea is to find the point on the curve that maximises both sensitivity and specificity. The CZ method simply finds the point where their product is maximised.

By considering the sqrt-log model of the upper 90% confidence interval of TDV versus sample size (Fig. 2) as equivalent to an ROC curve, we applied the CZ method to find the point on the curve where TDV and sample size are minimised. This is the point where we should achieve maximum value from the data. Further right along the curve, increasing the sample size would give a smaller improvement in trend reliability and is therefore not cost- or resource-effective. Since an ROC curve is intended for binary classification, the CZ method assumes that both sensitivity and specificity are on a 0-1 scale. TDV already ranges from 0-1, so we set sensitivity as 1 - TDV. We normalised sample size to a 0-1 scale by converting it to a proportion of the complete dataset (dividing by the total number of time series in the dataset). Since all datasets were the same size, the relationship between TDV and sample size was not altered by the conversion to a proportion. Specificity was then 1 - sample size. The optimal cut-point on the curve is defined as

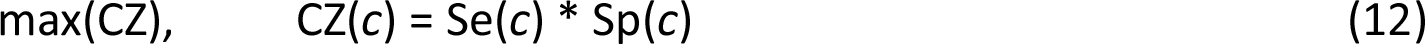

where Se is sensitivity, Sp is specificity, and *c* is any cut-point.

### 2.12. Minimum sample size for regional taxonomic groups

Minimum sample size was calculated by rearranging the formula for the multiple regression model to solve for sample size, and replacing the TDV variable in the formula with the cut-off value determined above. Values for the other variables in the formula were determined separately for each regional taxonomic group from the LPD, as follows: populations with fewer than two data points were removed, missing data was interpolated using the chain method (Collen et al., 2009), then the mean growth rate, *μ_pop_*, was calculated for each population. Growth rates were capped at [-1:1] before taking the mean, as in the LPI (McRae et al., 2017). Next, *μ_ds_*, *σ_ds_*, and *μ_ɳ_* were calculated. The mean time series length was calculated by dividing the total number of observations (after interpolation) by the total number of populations (excluding those with fewer than two data points). The calculated values were then placed into the model formula to determine minimum sample size.

### 2.13. Assigning reliability ratings to regional taxonomic groups

The actual number of populations in each regional taxonomic group was divided by the minimum sample size and multiplied by 100 to determine the percentage of the minimum sample size actually met by each group. Groups achieving 100% or greater were designated as reliable, those achieving between 50% and 100% were designated as data deficient, and those achieving less than 50% were designated as severely data deficient.

### 2.14. Correlations between reliability rating and LPI relative weighting

The Pearson’s product moment correlation coefficient test was performed to determine if there was any significant correlation between percentage of the minimum sample size achieved for each regional taxonomic group and the assigned relative weightings in the LPI for each group. The test was performed on the combined dataset as well as each individual system.

### 2.15. Modelling potential solutions

We used the model to simulate three different methods of improving trend reliability in the LPD: A) tracking unstudied populations for ten years, B) resampling previously-studied populations, and C) gathering more time series from existing studies. First, we generated 50 control datasets with a sample size of 200 and mean time series length of 14 (similar to the median values for regional taxonomic groups in the LPI of 180 and 13, respectively). We set *μ_ds_* to 0, *σ_ds_* to 0.25, and *μ_ɳ_* to 0.30. Using the same parameters, we then generated groups of 50 datasets with each of the following changes: group A had an extra 200 populations (total sample size: 400), but with observations only for the final ten years, to simulate tracking additional populations for ten years; group B had the final observation revealed on every sampled, degraded time series (total sample size: 200) to simulate resampling previously-studied populations; group C had an extra 200 randomly sampled populations (total sample size: 400) to simulate adding existing data to the LPD.

### 2.16. Coding and data

All trends for synthetic data were produced using original code designed to reproduce the functionality of the rlpi package (Freeman et al., 2021). All coding was done in R (R Core Team, 2021) using RStudio (RStudio Team, 2022). Fig. 1 was produced using Inkscape (Inkscape Project, 2020). All other figures were produced in R (R Core Team, 2021) using the ggplot2 package (Wickham, 2016). Population time series used to evaluate reliability of LPI trends are from the LPD (McRae et al., 2016). All original code is available on github at https://github.com/shawndove/DD_LPI.

## 3. Results

### 3.1. Regression model

The regression model contains five independent variables (Tables 1 and 2). Together they describe 62% of the variation (adjusted r-squared: 0.62) in the TDV associated with sampled trends, and with *F*(5, 29385) = 9,686, *p* < .001. All independent variables are statistically significant predictors, with p < 0.001. Interaction terms were also statistically significant but did not increase the adjusted r-squared of the model, so we left them out. RRMSEP is 0.231. Sample size is the most important variable affecting trend accuracy, with differences in importance between the other three variables comparatively small. Much of the unexplained variance from the model is due to random sampling. We confirmed this by remaking the model using the sample means, which resulted in an adjusted r-squared of 0.87. Using the square root of TDV instead of the log further increased the adjusted r-squared to 0.93. This was not the case for the model using the individual samples, where the log resulted in a higher adjusted r-squared than the square root.

**Table 2.** Multiple regression model of ln(TDV)

| coefficient | estimate | standard error | beta coefficient | t value | p value |
| --- | --- | --- | --- | --- | --- |
| (Intercept) | 3.957 | 0.04406 | – | 89.81 | < .001 |
| $\ln(\text{Sample size})$ | -0.8460 | 0.004441 | -0.6860 | -190.5 | < .001 |
| $\ln(\text{St. dev. of mean growth rate, } \sigma_{ds})$ | 0.7569 | 0.01630 | 0.1672 | 46.42 | < .001 |
| Mean growth rate, $\mu_{ds}$ | 8.057 | 0.1454 | 0.1989 | 55.42 | < .001 |
| Mean of population st. dev., $\mu_{\eta}$ | 1.503 | 0.02224 | 0.2426 | 67.57 | < .001 |
| Mean time series length | -0.03890 | 0.0007336 | -0.1917 | -53.02 | < .001 |

### 3.2. Maximum trend deviation value

Using the concordance probability method to select a cut point on the sqrt-log model of the 90% upper confidence interval of TDV versus sample size, we found a maximum TDV value of 0.176. After placing this value into the model equation and reorganizing to solve for sample size, we applied the model to the LPI to find the minimum number of populations needed for each regional taxonomic group.

### 3.3. Minimum sample size

The number of populations needed to achieve the TDV threshold for a reliable trend varies across taxonomic groups and realms (Table 3), but only weakly across systems, with medians of 269, 341, and 263 for terrestrial, freshwater, and marine systems, respectively. Fewer populations are needed in the global north (median: 213) than in the global south (median: 354). Birds show the highest variability, having both the smallest number of populations needed for any group (freshwater Nearctic birds: 19), and the largest (freshwater Afrotropical birds: 9,081; however, this value is an extreme outlier – see Discussion). Mammals have the smallest sample size requirements, with a median of 165, while fishes have the largest, with a median of 465. Reptiles and amphibians (combined) and birds fall in between, with medians of 274 and 286, respectively.

**Table 3.**
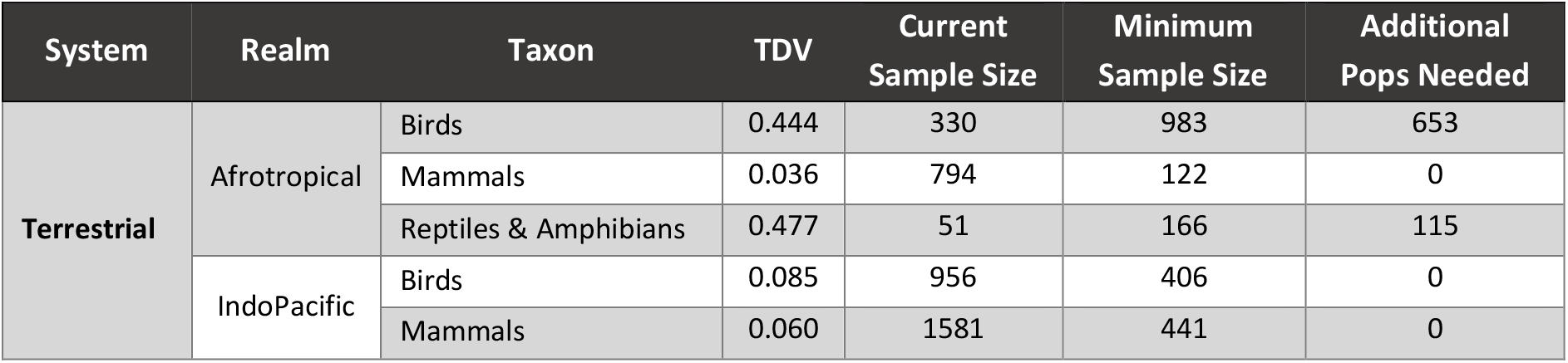
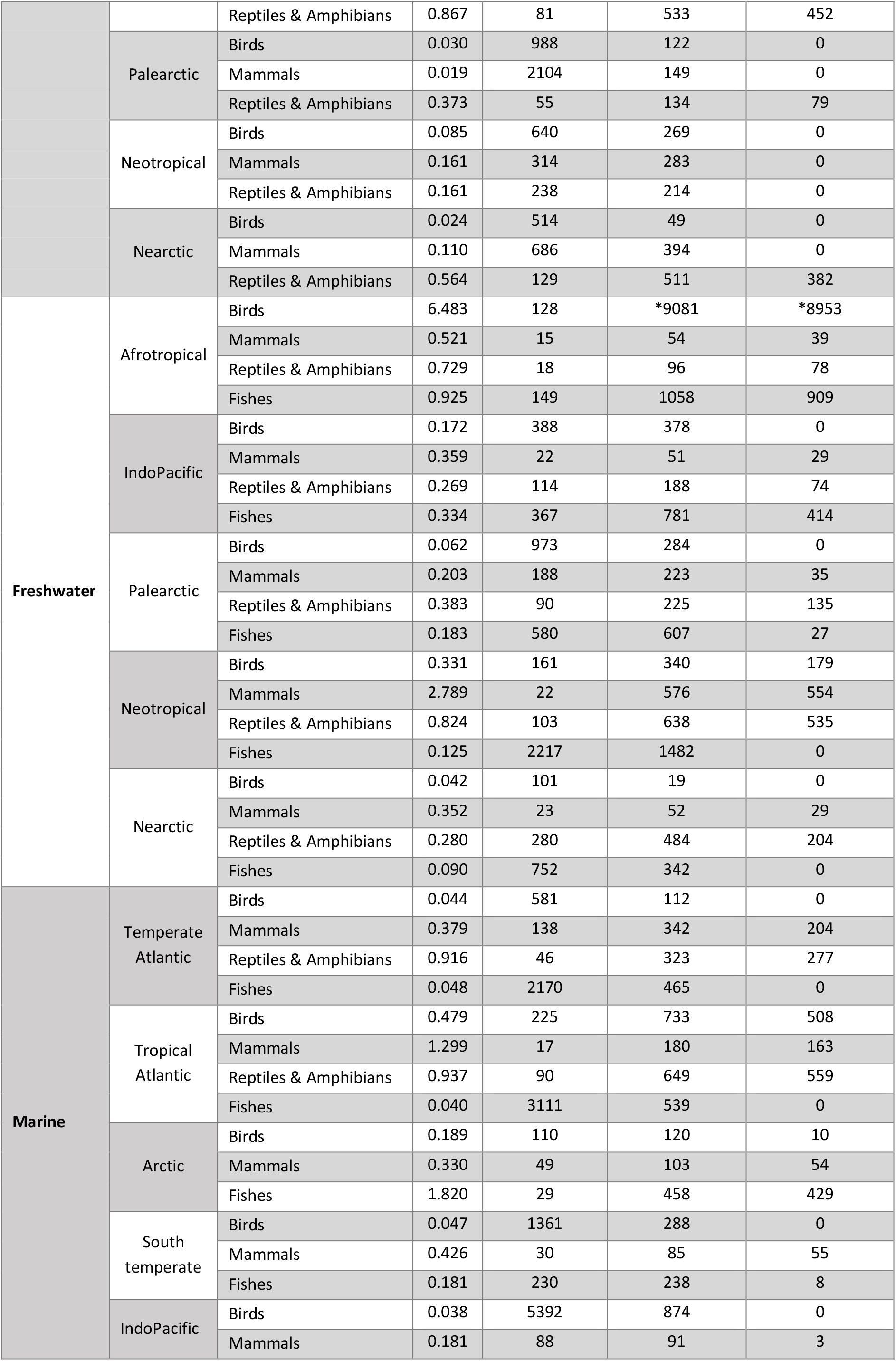
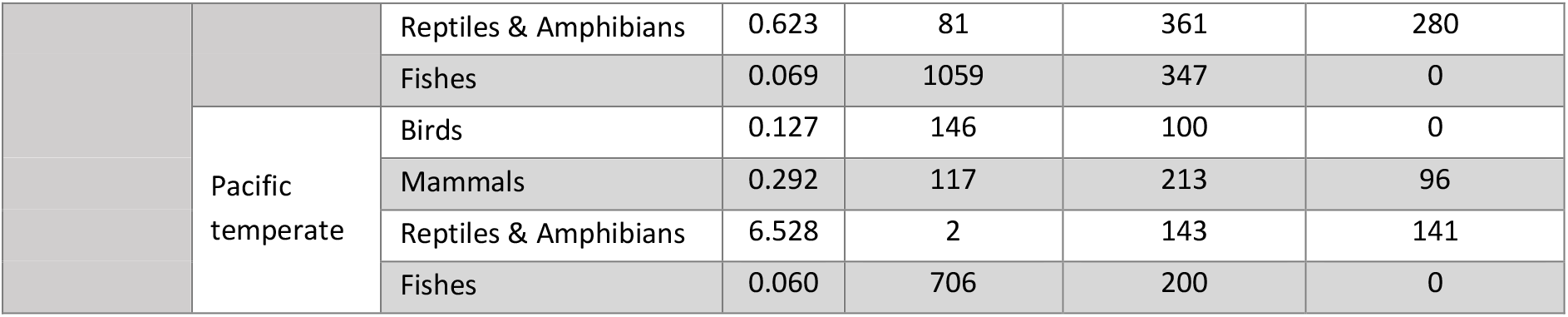
The trend deviation value, the current number of populations in the LPD, the minimum number of populations that would meet the reliability threshold, and the number of additional populations that must be added to achieve the reliability threshold for each regional taxonomic group in the LPD. Note that the trend deviation values here were calculated using the model formula and therefore occasionally fall outside of the 0-1 range of the Jaccard distance the TDV is based on. *Minimum sample size for freshwater Afrotropical birds is an extreme outlier. See Discussion for explanation.

| System | Realm | Taxon | TDV | Current Sample Size | Minimum Sample Size | Additional Pops Needed |
| --- | --- | --- | --- | --- | --- | --- |
| Terrestrial | Afrotropical | Birds | 0.444 | 330 | 983 | 653 |
|  |  | Mammals | 0.036 | 794 | 122 | 0 |
|  |  | Reptiles & Amphibians | 0.477 | 51 | 166 | 115 |
|  | IndoPacific | Birds | 0.085 | 956 | 406 | 0 |
|  |  | Mammals | 0.060 | 1581 | 441 | 0 |
|  |  | Reptiles & Amphibians | 0.867 | 81 | 533 | 452 |
|  | Palearctic | Birds | 0.030 | 988 | 122 | 0 |
|  |  | Mammals | 0.019 | 2104 | 149 | 0 |
|  |  | Reptiles & Amphibians | 0.373 | 55 | 134 | 79 |
|  | Neotropical | Birds | 0.085 | 640 | 269 | 0 |
|  |  | Mammals | 0.161 | 314 | 283 | 0 |
|  |  | Reptiles & Amphibians | 0.161 | 238 | 214 | 0 |
|  | Nearctic | Birds | 0.024 | 514 | 49 | 0 |
|  |  | Mammals | 0.110 | 686 | 394 | 0 |
|  |  | Reptiles & Amphibians | 0.564 | 129 | 511 | 382 |
| Freshwater | Afrotropical | Birds | 6.483 | 128 | *9081 | *8953 |
|  |  | Mammals | 0.521 | 15 | 54 | 39 |
|  |  | Reptiles & Amphibians | 0.729 | 18 | 96 | 78 |
|  |  | Fishes | 0.925 | 149 | 1058 | 909 |
|  | IndoPacific | Birds | 0.172 | 388 | 378 | 0 |
|  |  | Mammals | 0.359 | 22 | 51 | 29 |
|  |  | Reptiles & Amphibians | 0.269 | 114 | 188 | 74 |
|  |  | Fishes | 0.334 | 367 | 781 | 414 |
|  | Palearctic | Birds | 0.062 | 973 | 284 | 0 |
|  |  | Mammals | 0.203 | 188 | 223 | 35 |
|  |  | Reptiles & Amphibians | 0.383 | 90 | 225 | 135 |
|  |  | Fishes | 0.183 | 580 | 607 | 27 |
|  | Neotropical | Birds | 0.331 | 161 | 340 | 179 |
|  |  | Mammals | 2.789 | 22 | 576 | 554 |
|  |  | Reptiles & Amphibians | 0.824 | 103 | 638 | 535 |
|  |  | Fishes | 0.125 | 2217 | 1482 | 0 |
|  | Nearctic | Birds | 0.042 | 101 | 19 | 0 |
|  |  | Mammals | 0.352 | 23 | 52 | 29 |
|  |  | Reptiles & Amphibians | 0.280 | 280 | 484 | 204 |
|  |  | Fishes | 0.090 | 752 | 342 | 0 |
| Marine | Temperate Atlantic | Birds | 0.044 | 581 | 112 | 0 |
|  |  | Mammals | 0.379 | 138 | 342 | 204 |
|  |  | Reptiles & Amphibians | 0.916 | 46 | 323 | 277 |
|  |  | Fishes | 0.048 | 2170 | 465 | 0 |
|  | Tropical Atlantic | Birds | 0.479 | 225 | 733 | 508 |
|  |  | Mammals | 1.299 | 17 | 180 | 163 |
|  |  | Reptiles & Amphibians | 0.937 | 90 | 649 | 559 |
|  |  | Fishes | 0.040 | 3111 | 539 | 0 |
|  | Arctic | Birds | 0.189 | 110 | 120 | 10 |
|  |  | Mammals | 0.330 | 49 | 103 | 54 |
|  |  | Fishes | 1.820 | 29 | 458 | 429 |
|  | South temperate | Birds | 0.047 | 1361 | 288 | 0 |
|  |  | Mammals | 0.426 | 30 | 85 | 55 |
|  |  | Fishes | 0.181 | 230 | 238 | 8 |
|  | IndoPacific | Birds | 0.038 | 5392 | 874 | 0 |
|  |  | Mammals | 0.181 | 88 | 91 | 3 |
|  |  | Reptiles & Amphibians | 0.623 | 81 | 361 | 280 |
|  |  | Fishes | 0.069 | 1059 | 347 | 0 |
|  | Pacific<br>temperate | Birds | 0.127 | 146 | 100 | 0 |
|  |  | Mammals | 0.292 | 117 | 213 | 96 |
|  |  | Reptiles & Amphibians | 6.528 | 2 | 143 | 141 |
|  |  | Fishes | 0.060 | 706 | 200 | 0 |

### 3.4. Trend reliability

Reliability varies strongly across realms, taxonomic groups, and systems (Figs 3 & 4). Terrestrial trends are the most reliable and freshwater trends the least. Terrestrial and freshwater trends are more reliable in the global north than in the global south, except for terrestrial reptiles and amphibians. Marine bird trends are more reliable in temperate areas than the tropics, while marine fish trends are more reliable in tropical waters than polar. Globally, bird trends are the most reliable, but are nonetheless poor in the tropics, especially Africa. Reptile and amphibian trends are data deficient everywhere except the terrestrial Neotropical realm, and marine and freshwater mammal trends are data deficient everywhere (although marine IndoPacific mammals are very close to the threshold at 97%).

**Fig. 3.**
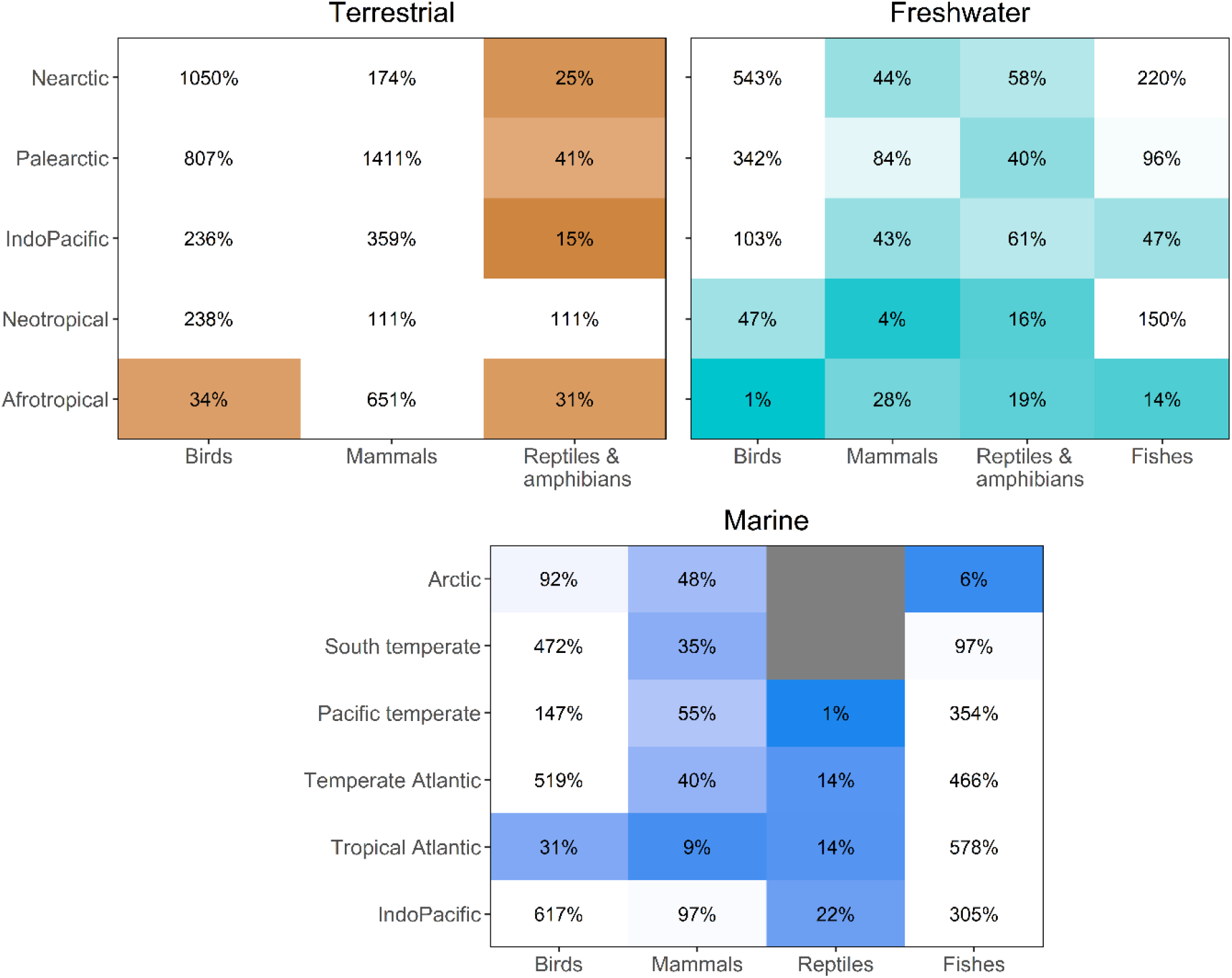
Proportion of the total amount of time series data needed to achieve the trend reliability threshold that each regional taxonomic group in the LPD currently contains. A score of 100% or greater means that group already has enough data to produce a reliable trend. A white box refers to a group that meets the reliability threshold, while a coloured box means the threshold has not been met. The further the group is from meeting the threshold, the more intense the colour. A grey box refers either to a group that could not be evaluated because there was too little data (South temperate marine reptiles) or due to an invalid realm-taxon combination (there are no marine reptiles in the Arctic).

**Fig. 4.**
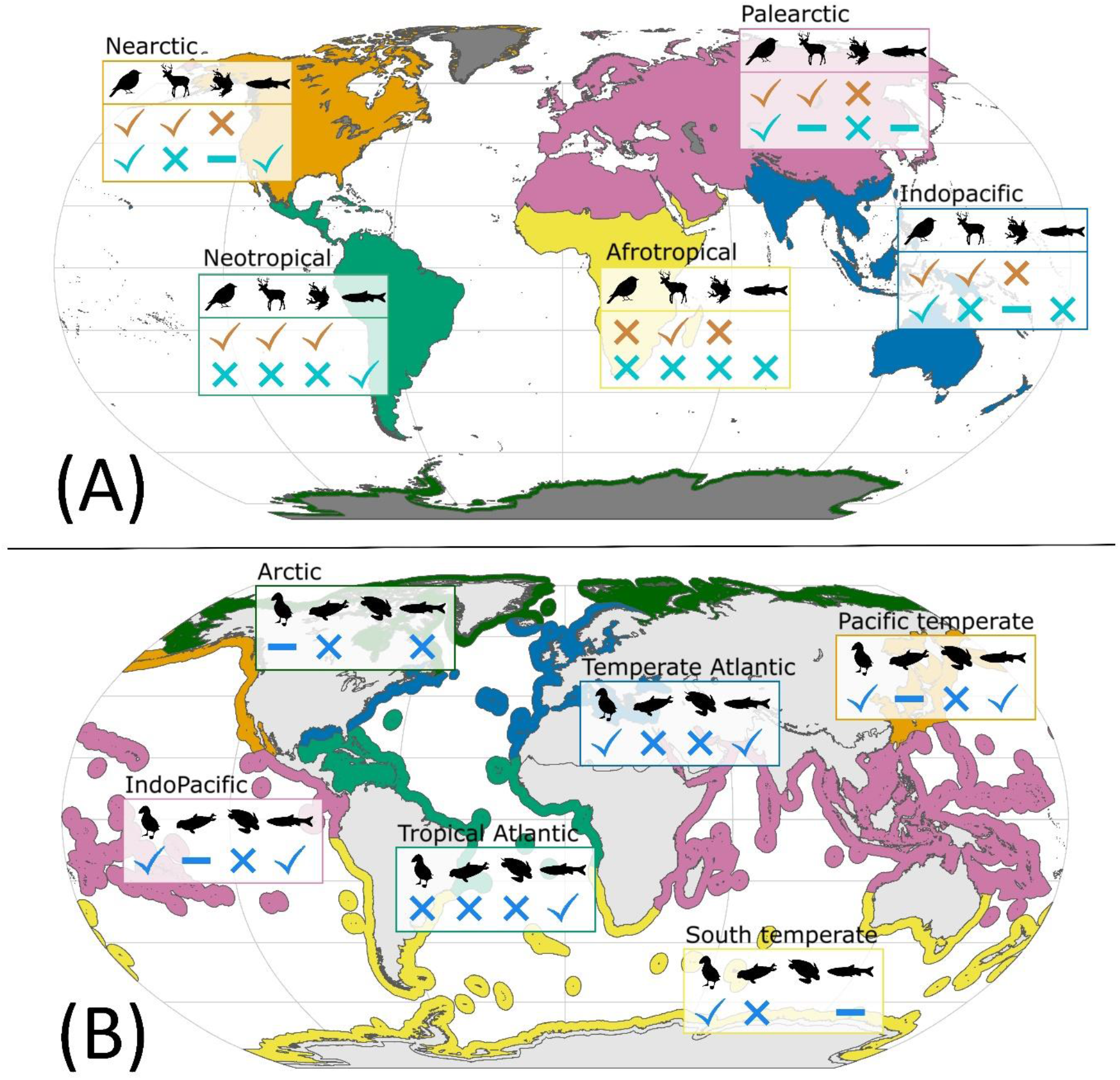
Reliability of regional taxonomic group trends in the LPI, grouped by system, realm, and taxon. Map A shows the terrestrial (top) and freshwater (bottom) results. Map B shows the marine results. Reliability scores are binned into three categories, according to the number of time series in the LPD relative to the minimum sample size needed to achieve the TDV threshold. A check mark means that group has at least 100% of the minimum sample size and is considered reliable, a dash means it is data deficient (50-99%), and an X mark means it is severely data deficient (< 50%).

The regional taxonomic groups with the greatest potential to affect the reliability of aggregated LPI trends are exclusively tropical (Fig. 5), due to a combination of high relative weighting and low reliability scores. The eight groups of greatest concern include five freshwater and three terrestrial groups, but no marine groups. All are from the tropics. Fishes, birds, and reptiles and amphibians are represented, with mammals absent. Overall, the reliability scores of regional taxonomic groups did not show a statistically significant correlation with their relative weightings in the LPI, *r*(55) = 0.085, *t* = 0.64, *p* = 0.53. Likewise, there were no statistically significant correlations for individual systems, with terrestrial *r*(13) = -0.40, *t* = -1.56, *p* = 0.14; freshwater *r*(18) = -0.09, *t* = -0.39, *p* = 0.70; and marine *r*(20) = 0.40, *t* = 1.95, *p* = 0.066.

**Fig. 5.**
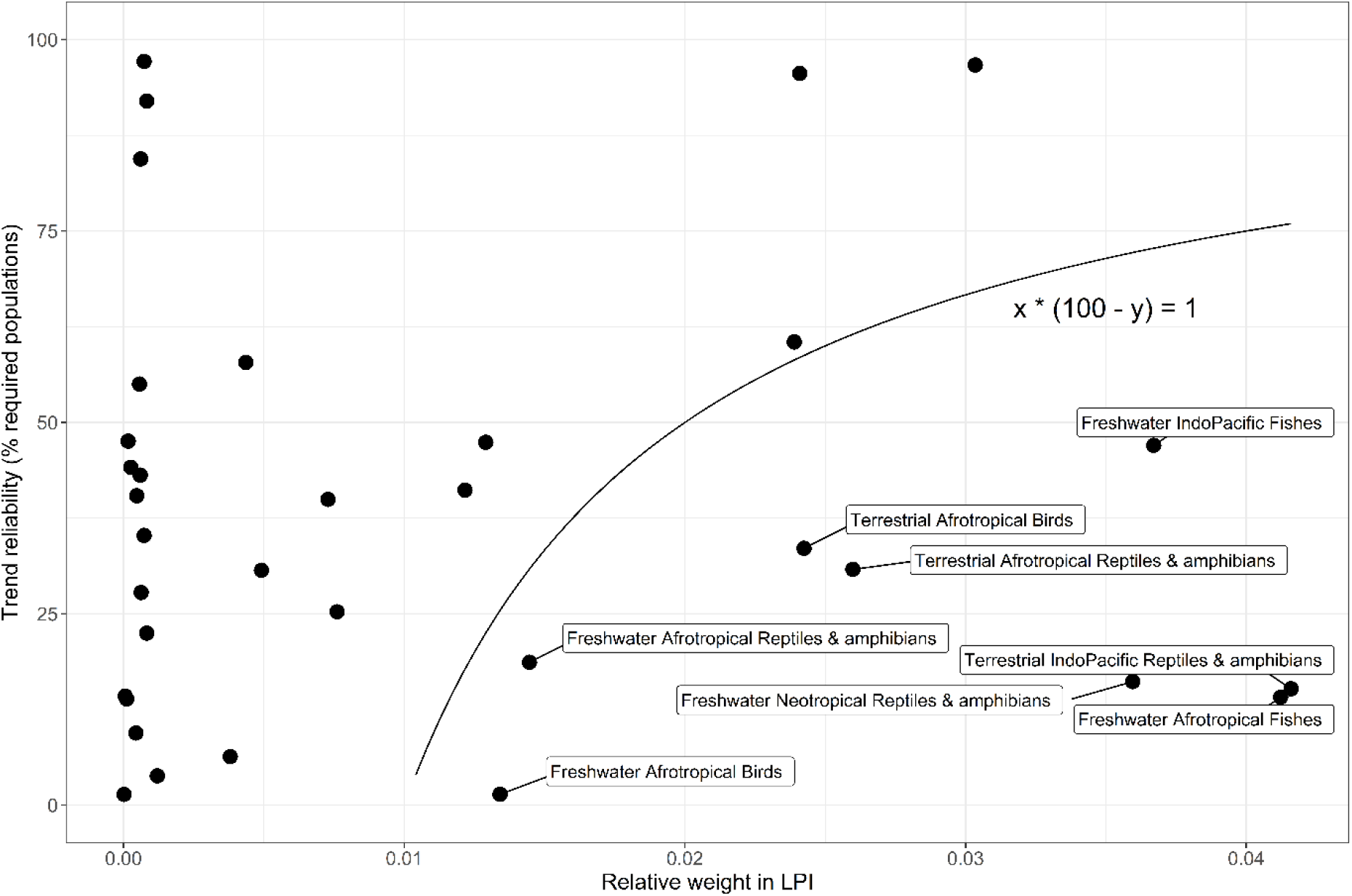
Trend reliability of regional taxonomic groups in the LPD (measured as the percentage of populations in the LPD relative to the number required to achieve the TDV threshold) versus the relative weighting applied to each group when calculating aggregated LPI trends. Only groups with reliability ratings below the threshold (less than 100%) are included here. To determine the groups having the strongest negative effect on the reliability of aggregated LPI trends, we calculated relative weight * (100 – reliability) and labelled the groups with a value higher than 1.

### 3.5. Modelling potential solutions

Revealing the final year observation (equivalent to resampling previously-studied populations) for every population improved the median TDV by 6.5%, while adding 200 additional time series to the sample with observations only in the final ten years (equivalent to tracking 200 unstudied populations for ten years) improved the mean TDV by 11% (Fig. 7). By contrast, simply doubling the sample size (equivalent to randomly adding 200 existing time series to the LPD) improved the median TDV by 43%. This solution showed a statistically significant improvement in trend accuracy compared to the control group (p < 0.001).

**Fig. 7.**
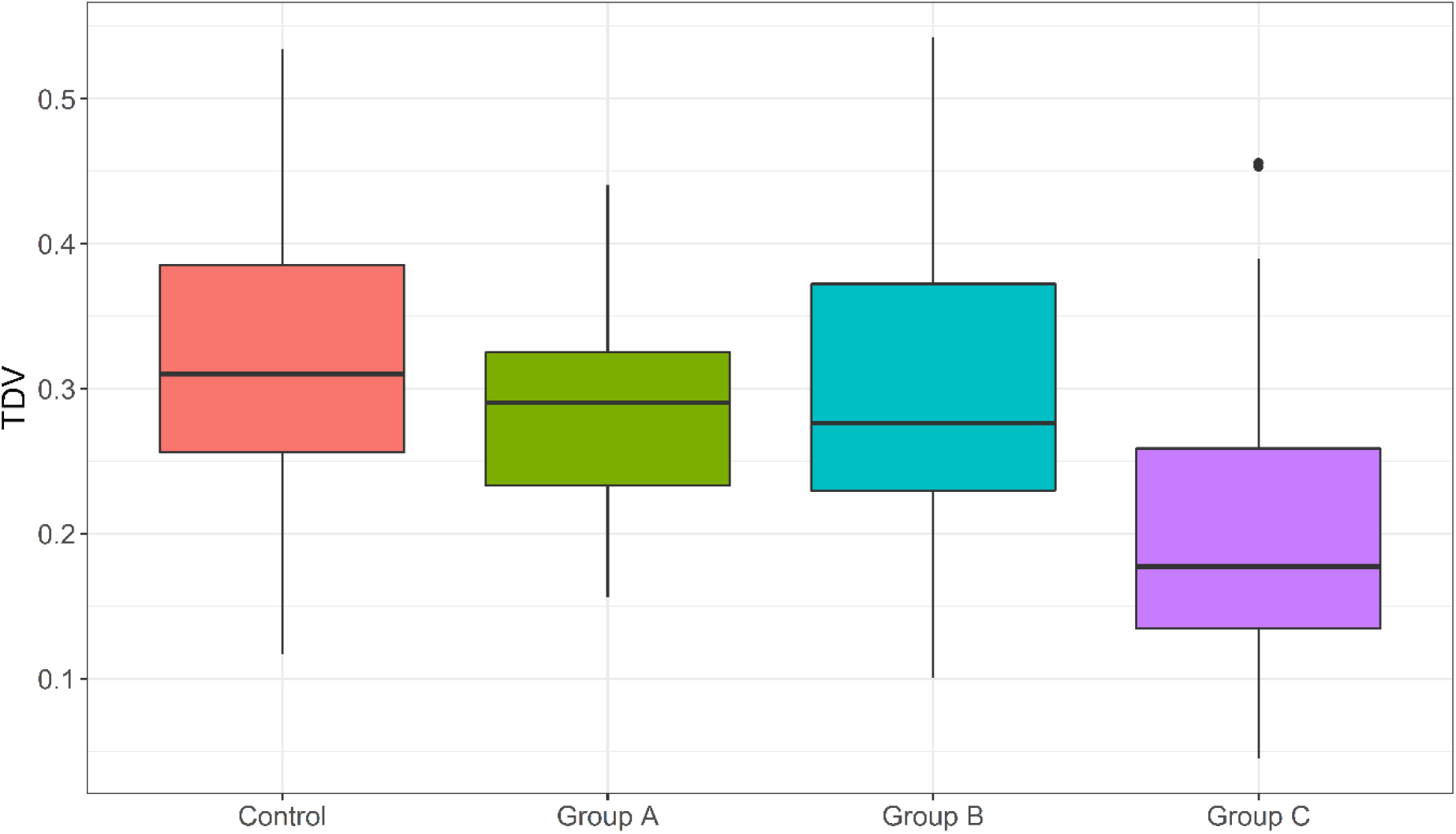
The effect on trend accuracy of potential solutions to data deficiency in LPI regional taxonomic groups. The control group has a sample size of 200 and mean time series length of 14. Group A has an additional 200 time series with observations only in the final ten years of the index to simulate a ten-year data blitz. In group B, the final observation has been added back in for every time series to simulate resampling of previously-studied populations. Group C is like the control group, but the sample size has been doubled to 400 to simulate adding additional pre-existing studies to the LPI.

## 4. Discussion

Understanding the changing global state of biodiversity is crucial to making good policy and conservation decisions and ‘bend the curve’ of biodiversity loss (Mace et al., 2018). Acquiring accurate and comprehensive data is crucial, but the first step is to answer the question: what do we actually know? The present study quantifies the reliability of trends for each regional taxonomic group in the Living Planet Index and estimates the number of population time series needed to meet a standard of expected accuracy.

We used synthetic population time series datasets to construct a multiple regression model of trend accuracy by comparing trends of degraded samples with the trends of the full, undegraded datasets using a distance measure (Fig. 1). We applied the model to regional taxonomic groups in the Living Planet Database to reveal that the majority need additional data for their trends to be considered reliable. Data deficiency is a problem globally, but is more pronounced in the tropics. This is consistent with the analysis of geographical representativeness in McRae et al. (2017), which tested proportional representativeness of biodiversity compared to the global dataset and found that species groups in tropical realms are underrepresented. Bird trends are the most reliable and reptiles and amphibians the least. This is consistent with the picture of species representation in the LPD presented in McRae et al. (2017) and is unsurprising given that monitoring and data collection for birds is more extensive than for reptiles and amphibians (Oliver et al., 2021; Scheele et al., 2019), especially with the rise of citizen science (Oliver et al., 2021). However, many of our reliability scores differ from what would be expected given McRae et al. (2017)’s analysis of taxonomic representativeness. McRae et al. (2017) found that all Nearctic taxonomic groups are overrepresented, yet in our analysis Nearctic terrestrial and freshwater reptiles and amphibians as well as Nearctic freshwater mammals score as data deficient. The starkest differences occur in the marine system, where mammals and marine reptiles are overrepresented by species in all realms (except South temperate reptiles, which are not represented at all) but which we found to be data deficient in all realms. In contrast, marine fishes are underrepresented by species numbers (McRae et al., 2017), but we found that in all except the Arctic realm marine fishes are data-rich enough to produce reliable trends. These results strongly suggest that the percentage of species represented does not tell the whole story.

Geographical and taxonomic biases in the distribution of data in the LPI are well-known (McRae et al., 2017), and reflect underlying biases in the availability of data (Boakes et al., 2010; Collen et al., 2008; Yesson et al., 2007). In 2017, McRae et al. introduced a weighting system to the LPI, which accounts for the estimated number of species in each regional taxonomic group to reduce representational bias. One problem with this is that most of the world’s vertebrate species are located in the tropics (Collen et al., 2008; McRae et al., 2017), which are underrepresented in the LPD (McRae et al., 2017). Our concern was that if trends from these areas are the least reliable due to data deficiency, then the LPI could have simply replaced one problem, representation bias, with another: overreliance on data deficient trends. Indeed, our analysis shows that all regional taxonomic groups with a high relative weight and low reliability rating (bottom right of Fig. 5) are tropical. Surprisingly, though, we did not find a statistically significant negative correlation between reliability of trends and their relative weights in the LPI. This also holds true for the terrestrial and freshwater systems when considered separately (the marine system actually shows a positive correlation), and is consistent with Nori et al. (2020), who found that species richness and knowledge gaps are not always correlated.

According to our model, the size of a dataset, i.e., the number of species or populations existing in the real world for any regional taxonomic group, is unimportant to the calculation of trend reliability for a given sample, as long as the sample represents less than half of the time series in the dataset (Fig. S1). In other words, it is the absolute number of populations represented in the sample that matters, regardless of whether that sample represents 1% or 50% of the total populations in a regional taxonomic group. There are two principles working to cause this seemingly counterintuitive effect. First, the relationship between population size and the sample size needed to reach a desired level of precision is logarithmic and becomes more extreme at lower levels of precision (Israel, 1992). This means that a small sample size should be able to estimate a large population almost as well as it can estimate a small population. Second, there are limitations to the level of trend accuracy that can be achieved, regardless of sample size, because most time series in our simulated samples (and in the LPD) are much shorter than the length of the trend being estimated. Short time series tend to produce more extreme trends (Leung et al., 2020) and are less likely to accurately reflect long-term trends for individual populations (Wauchope et al., 2019). They also reduce the number of observations used for the calculation of group trends. For example, even if the mean time series length was 50% of the length of a trend (mean time series lengths for all regional taxonomic groups in the LPD are much shorter than that), if those time series were randomly distributed in time, only about 4% of them would begin at the first year and about 4% would end at the final year. Thus, the crucial early and final years of the trend would depend on only a fraction of the observations that the sample size indicates. This randomised distribution of time series across the trend results in less accurate trends than would be possible if observations were evenly distributed across time points (confirmed through simulations – see Fig. S5). This issue is slightly complicated in the LPD. On one hand, the database begins 20 years earlier than the index, giving time for the number of observations to increase before measuring the trend. But on the other hand, there is a delay in getting recent studies into the LPD (McRae et al., 2017), reducing the number of observations in the final years even more than a random distribution would suggest (see Fig. S6).

This dramatic fall-off of observations suggests that more data is needed for the LPI to reliably reflect changes in the status of global vertebrate biodiversity over the past decade. While a reduction in the delay involved in getting new studies into the LPD might help, increasing the number of populations in the LPD is only possible to the extent that the necessary data exists. Therefore, we simulated two potential ways of generating new data to improve trend reliability: A) a global data blitz, with researchers coordinating to track as many unstudied populations as possible for ten years to generate new time series, and B) resampling already-studied populations to uncover recent changes and lengthen existing time series (Fig. 7). Both solutions had a slight but non-significant positive effect on trend accuracy, but were far less effective than adding existing data (as is currently done for the LPD). It is likely that both solutions have a greater effect on the accuracy of the final portion of the trend than on the overall trend, but further study would be required to be certain. Either way, resampling would be more efficient than a data blitz, as the same improvement could be achieved in one year instead of ten. In the long term, tracking additional populations is essential to completing our picture of biodiversity change. However, natural stochasticity means that short time series are of limited value in generating reliable trends (Wauchope et al., 2019), so tracking additional populations takes time to pay dividends.

There is another limitation underlying the LPI, which cannot be solved by generating new data. All trends in the LPI begin in the year 1970, which is set as the base year for calculating the index values. Past trends can only be determined by existing data; therefore, while there may be some currently inaccessible data that either could be shared or made available for confidential storage in the LPD (Saha et al., 2018), there are likely to be severe limitations to relieving data deficiency for the early years of the LPI. However, other potential solutions could be examined in future studies. One would be to begin the index at a later year in which there is more data available (e.g., 1990). Another would be to change the base year for calculating the index to a more data-rich year, thus increasing the uncertainty around the early years of LPI trends (Gregory et al., 2019). The downside is that the interpretation of trends would be different. The LPI would no longer measure change in global vertebrate biodiversity relative to 1970, but relative to another year. Much of the change currently recorded in the index would have already occurred before the base year. Another potential solution would be to use other kinds of data, such as log books and catch records (e.g. Josephson et al., 2008), genetics (e.g. Beaumont, 2003), trade records (e.g. Collins et al., 2020), and land use/climate change modelling (e.g. Visconti et al., 2015) to infer historical abundance estimates for populations where no monitoring took place.

Our modelling approach to quantifying trend reliability is subject to several limitations. Certain aspects of the underlying data, such as the distribution of observations and biases in which populations or species are tracked, are too complex to be included as factors in the model, but nonetheless may play a significant role in determining trend reliability. For example, monitoring efforts tend to focus on species at higher risk of extinction (Scheele et al., 2019). Many amphibian populations in the LPD were tracked because they were declining due to the devastating disease *chytridiomycosis*. This could negatively bias trends and falsely reduce variance in growth rates, leading the model to overestimate reliability because it assumes that tracked populations are randomly selected. On the other hand, Murali et al. (2022) found that population coverage in the LPD is biased towards protected areas, where species are less likely to be threatened, therefore potentially causing a positive bias in LPI trends. Another common phenomenon in the LPD is that time series are non-randomly distributed across time and/or space. For example, while some biodiversity hotspots (e.g., tropical Africa) are poorly known, others, especially islands (e.g., Madagascar), are well-studied (Nori et al., 2020), and this may bias entire realms. In the Afrotropical realm, six (12%) of the 51 terrestrial reptile and amphibian time series in the LPD are from Round Island (a tiny uninhabited island near Mauritius) and more than half (57%; 29/51) are from a single study that took place at a reserve in Madagascar over a nine-year period; only seven (14%) are from mainland Africa, and of the seven, four are from a single study at a reserve in Nigeria. In this case, the model likely severely underestimates the amount of data needed to get a reliable trend. While this is an extreme example, it shows that there are important underlying aspects of the data that cannot be assessed by a model. Fortunately, these issues tend to diminish when more data is present, and thus should not have a large effect on trends assessed as reliable.

The model also assumes that adding additional time series to the LPD will maintain the parameters of the regional taxonomic group to which they are added (e.g., the mean time series length and the level of variance in population mean growth rates will not change). This can result in the model severely overestimating the numbers of populations required to achieve a reliable trend. For example, it suggested that 9,087 populations of freshwater Afrotropical birds are needed. This likely occurred due to problems with the existing data. Although there are 128 freshwater Afrotropical bird populations in the LPD, most of them are short and/or sporadically observed (the mean number of observations is 4.0), and observations are clustered in the 1990s and 2000s, with only a single time series containing observations after 2009. However, this is an issue only for small or exceptionally poor quality samples (e.g. short time series, few studies, biased distribution in time and space), and if more and better time series are added to the LPD, the model should improve its estimates.

Another limitation of our modelling approach is that we could not correct for the sizes of the ‘real-world’ datasets (the number of populations that exist) that the LPD ‘samples’ are drawing from, and therefore may overestimate the sample size needed to achieve a reliable trend for very small datasets. Although there are estimates of the number of species for each regional taxonomic group, our model uses populations as the base unit to measure sample size. We chose to base sample size on populations rather than species for two reasons. First, we found that mean growth rates within the LPD vary almost as much between populations within a species as they do between species. Therefore, we cannot assume that the trend of a population represents the trend of the species it belongs to any better than it represents the trend of its entire regional taxonomic group. Second, localised threats such as land-use change and habitat destruction are likely to affect some populations within a species disproportionately. Population extinctions also occur much more frequently than species extinctions, and may serve as a prelude (Ceballos et al., 2017). However, a population is not a well-defined unit, and we do not have estimates of how many populations each species or regional taxonomic group is composed of. While our testing suggested we can assume the number of existing populations to be unimportant in determining trend reliability, this assumption breaks down when the sample comprises a large percentage of the dataset. It is unlikely that any regional taxonomic groups currently approach this level of representation within the LPD, but it is nonetheless an important caveat to be aware of.

Despite these caveats, the results of our study reveal the strengths and weaknesses in our understanding of global vertebrate biodiversity, highlighting the regional taxonomic groups for which we have enough data to make responsible decisions, as well as those on which future data gathering and collation efforts should focus. Some underlying aspects of the data create biases that are not taken into account by our modelling approach, and more fine-scale studies on gaps in population trends should be performed to better understand these biases and where to divert scientific resources. We show that revisiting previously-studied populations is a quick and efficient way to improve trend reliability for data deficient groups until more long-term studies can be completed and made available. The modelling approach we use to quantify trend reliability can also be generalized to assess other global and/or regional biodiversity indices that utilize population time series data. We are facing an urgent global biodiversity crisis made worse by biased and deficient data, but through careful study and cooperative global efforts we can solve the data problem and begin to ‘bend the curve’ of biodiversity toward a positive trend.

## Acknowledgements

We thank Sean Jellesmark, Gonzalo Albaladejo-Robles, and Bouwe Reijenga for their support. This project has received funding from the European Union’s Horizon 2020 research and innovation programme under the Marie Skłodowska-Curie grant agreement No 766417.

