## Supplementary Figures for "How much data do we need? Reliability and data deficiency in global vertebrate biodiversity trends"

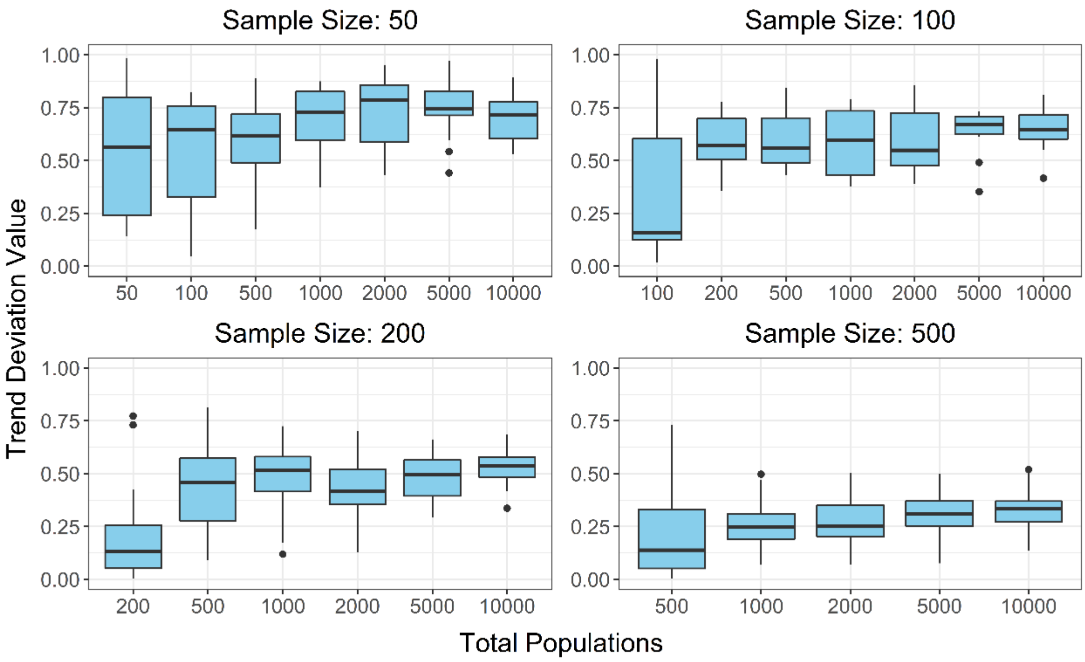


Fig. S1. Trend deviation value (TDV) vs total number of populations in dataset, at different sample sizes. All other parameters are fixed – $\mu_{ds}$: 0; $\sigma_{ds}$: 0.4; $\mu_{\eta}$: 0.6; populations per species: 20; mean time series length: 20; trend length: 50; $\mu_{\varepsilon}$: 0; $\sigma_{\varepsilon}$: 0. Each box represents the mean values of 10 datasets, with 20 samples per dataset. There is no clear effect of dataset size on TDV. While TDV is lower when the dataset is equal to the sample size (TDV is non-zero due to degradation of the samples), there is no clear trend in median TDVs as the number of populations in the dataset increases from twice the size of the sample to 10,000 populations.

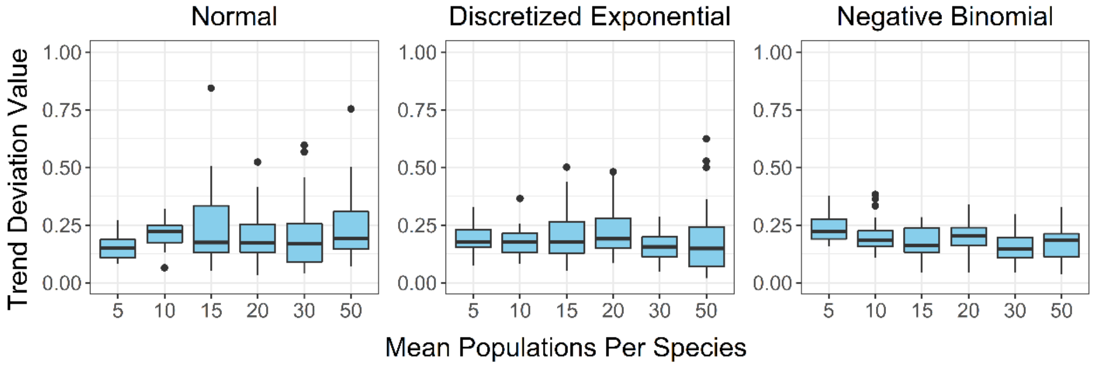


Fig. S2. Trend deviation value vs mean number of populations assigned to each species, using three different distributions: a normal distribution; an exponential distribution discretized by rounding to the nearest whole number; and a negative binomial distribution with zeros removed by adding 1 to every value (the resulting increase of the mean was accounted for). All other parameters are fixed – dataset size: 1000; sample size: 200; $\mu_{ds}$: 0; $\sigma_{ds}$: 0.2; $\mu_{\eta}$: 0.2; mean time series length: 20; trend length: 50; $\mu_{\varepsilon}$: 0.15; $\sigma_{\varepsilon}$: 0.1. Each box represents the mean values of 20 datasets, with 20 samples per dataset. Neither the mean number of populations assigned to each species, nor the distribution used to assign them, shows any effect on trend accuracy.

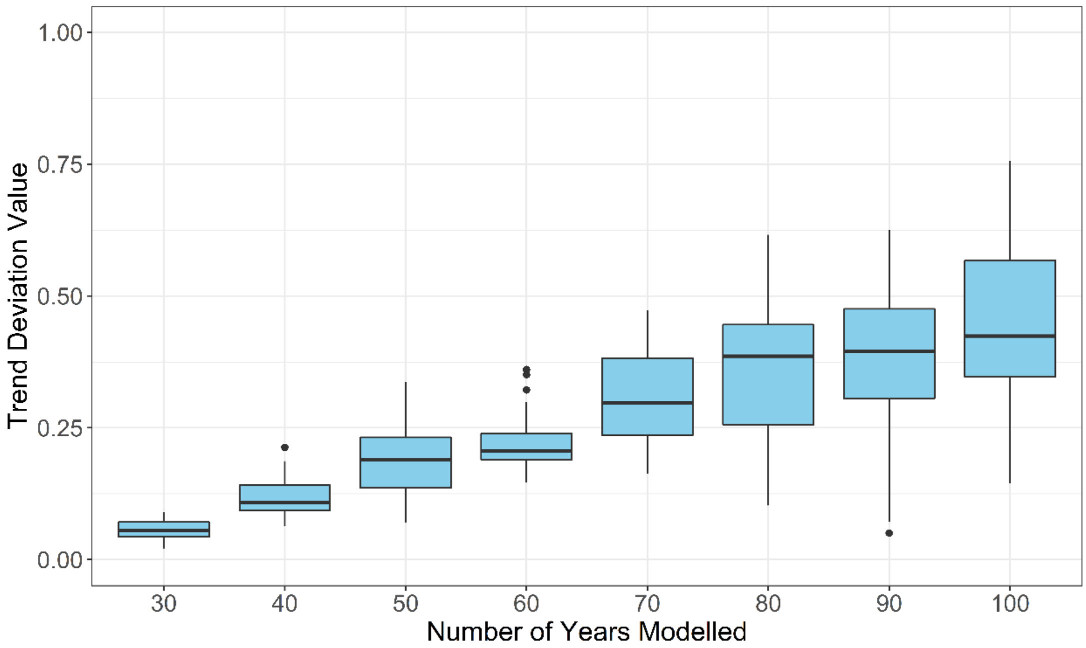


Fig. S3. Trend deviation value vs length of trend. All other parameters are fixed – dataset size: 1000; sample size: 200; $\mu_{ds}$: 0; $\sigma_{ds}$: 0.2; $\mu_{\eta}$: 0.2; populations per species: 20; mean time series length: 20; $\mu_{\varepsilon}$: 0.15; $\sigma_{\varepsilon}$: 0.1. Each box represents the mean values of 20 datasets, with 20 samples per dataset. The median and the range of the TDV increase as trend length increases; this likely occurs because the mean time series length and sample size are static, resulting in fewer observations per year.

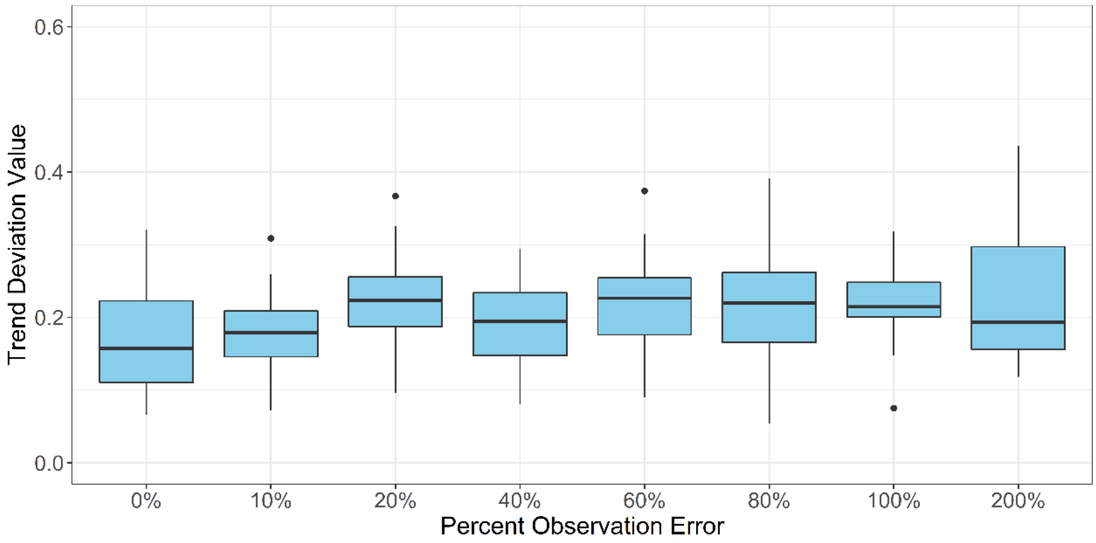


Fig. S4. Trend deviation value vs percent observation error. All other parameters are fixed – dataset size: 1000; sample size: 200; $\mu_{ds}$: 0; $\sigma_{ds}$: 0.2; $\mu_{\eta}$: 0.2; populations per species: 20; mean time series length: 20; trend length: 50. Each box represents the mean values of 20 datasets, with 20 samples per dataset. The percentage of observation error has no effect on trend accuracy.

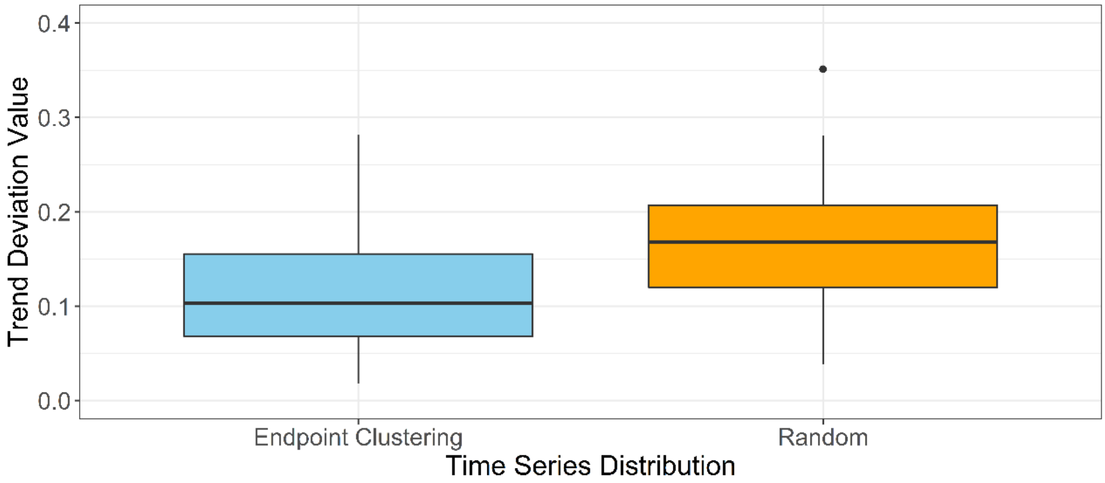


Fig. S5. Trend deviation value vs time series distribution. Datasets assigned to "Endpoint Clustering" had time series randomly assigned to begin at the first year of the trend or at end at the final year of the trend, while datasets assigned to "Random" had time series randomly distributed across the simulated years of the trend. All other parameters are fixed – dataset size: 1000; sample size: 200; $\mu_{ds}$: 0; $\sigma_{ds}$: 0.2; $\mu_{\eta}$: 0.2; populations per species: 20; mean time series length: 25; trend length: 50; $\mu_{\varepsilon}$: 0.15; $\sigma_{\varepsilon}$: 0.1. Each box represents the mean values of 40 datasets, with 20 samples per dataset.

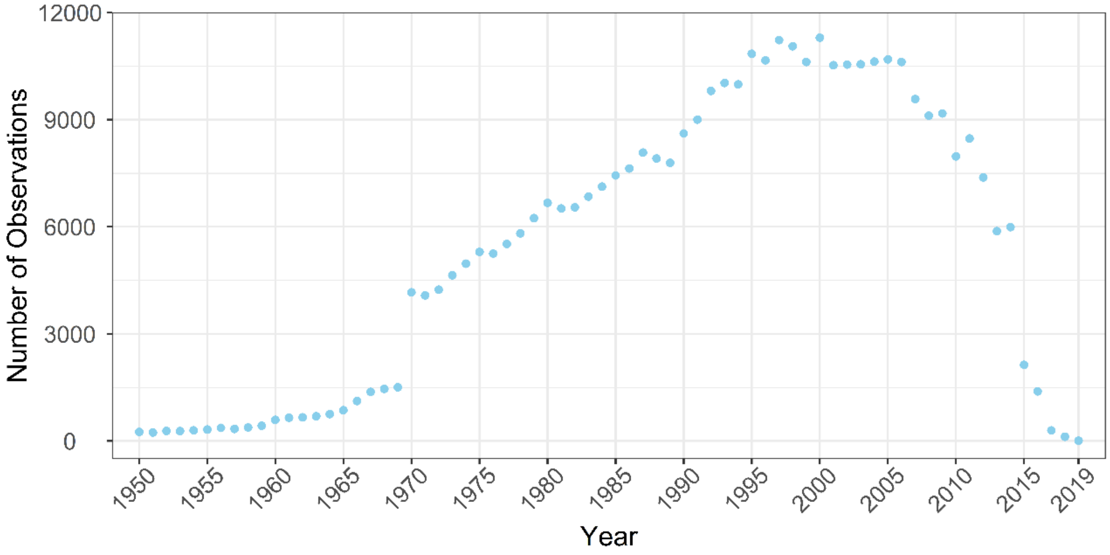


Fig. S6. Total number of observations recorded in the Living Planet Database for each year from the start of the database until 2019 (the most recent observations in the version of the database we used for our analysis). There are 250 observations for the year 1950 and 9 observations for the year 2019. The year 2000 has the highest number of observations, at 11,297.
